# Efficacy and Specificity of Melanopsin Reporters for Retinal Ganglion Cells

**DOI:** 10.1101/2022.04.26.489554

**Authors:** Ryan Maloney, Lauren Quattrochi, James Yoon, Rachel Souza, David Berson

**Author notes:** Department of Neurological Surgery, University of Pittsburgh Medical Center, Pittsburgh, PA, USA.

## Abstract

Intrinsically photosensitive retinal ganglion cells (ipRGCs) are specialized retinal output neurons that mediate behavioral, neuroendocrine, and developmental responses to environmental light. There are diverse molecular strategies for marking ipRGCs, especially in mice, making them among the best characterized retinal ganglion cells. With the development of more sensitive reporters, new subtypes of ipRGCs have emerged. We therefore tested high-sensitivity reporter systems to see whether we could reveal yet more. Substantial confusion remains about which of the available methods, if any, label all and only ipRGCs. Here, we compared many different methods for labeling of ipRGCs, including anti-melanopsin immunofluorescence, Opn4-GFP BAC transgenic mice, and Opn4^cre^ mice crossed with three different Cre-specific reporters (Z/EG; Ai9; and Ai14) or injected with Cre-dependent (DIO) AAV2. We show that Opn4^cre^ mice, when crossed with sensitive Cre-reporter mice, label numerous ganglion cell types that lack intrinsic photosensitivity. Though other methods label ipRGCs specifically, they do not label the entire population of ipRGCs. We conclude that no existing method labels all and only ipRGCs. We assess the appropriateness of each reporter for particular applications and integrate findings across reporters to estimate that the overall abundance of ipRGCs among mouse retinal ganglion cells may approach 11%.

## Introduction

The mouse retina has at least 40 types of retinal ganglion cells (RGCs)(Sanes and Masland, 2015; Baden et al., 2016; Bae et al., 2018; Tran et al., 2019; Goetz et al., 2022). Each type distills a distinctive form of visual information from the retinal image and conveys it to specific brain targets. The steady emergence of new and powerful genetically modified mouse models for tagging and manipulating specific RGC types (Hattar et al., 2002; Huberman et al., 2008; Martersteck et al., 2017) has driven a dramatic growth in our understanding of this diversity, including detailed descriptions of the structure, function, and brain connectivity of diverse RGC types and the roles they play in visual guided behavior.

Intrinsically photosensitive retinal ganglion cells (ipRGCs) are unique among RGCs in expressing melanopsin, which enables them to respond to light independently from rods and cones. Techniques exploiting the melanopsin gene to label ipRGCs have revealed at least six subtypes of ipRGCs, each with characteristic morphology, stratification, response properties, and brain projections (Berson et al., 2010; Ecker et al., 2010; Estevez et al., 2012; Stabio et al., 2017; Quattrochi et al., 2018).

The original and simplest method of identifying ipRGCs is anti-melanopsin immunostaining (Provencio et al., 2000). Because melanopsin protein is distributed throughout the plasma membrane, immunolabeling reveals somadendritic and even axonal morphology of many ipRGCs. This reveals three ipRGC subtypes each with a unique pattern of dendritic stratification: the M1, M2 and M3 cells (Berson et al., 2010). An additional advantage of the immunolabeling approach is that it is effective in diverse species that lack the powerful molecular reagents available in mice. However, a limitation of the immunostaining approach is that it doesn’t reliably reveal ipRGC subtypes that express relatively low levels of melanopsin (in mice, the M4, M5, and M6 types), despite their demonstrable autonomous, melanopsin-dependent photosensitivity (Estevez et al., 2012; Stabio et al., 2017; Quattrochi et al., 2018).

ipRGCs have also been revealed in mice modified genetically to report melanopsin expression. The first to be developed (Hattar et al., 2002) knocked in *tau-lacZ* into the melanopsin locus; cells expressing the reporter were revealed by enzyme histochemistry. It now appears that this approach labels only the M1 type of ipRGC, which expresses more melanopsin than other ipRGC types (Hattar et al., 2006). A second approach has been to generate transgenic animals that express fluorescent proteins under the control of the melanopsin promotor through introduction of bacterial artificial chromosomes (BACs) (Schmidt et al., 2008; Do et al., 2009) In these mice too, the level of expression of the reporter appears linked to the level of melanopsin expression because only the heavily melanopsin-immunopositive ipRGC types are tagged (i.e., the M1, M2 and M3 subtypes) (Schmidt et al., 2008).

Yet another strategy to label ipRGCs uses the melanopsin promoter to drive expression of Cre recombinase. When coupled with an appropriate reporter mouse or virus, Cre yoked to *opn4* transcription triggers expression of a fluorescent protein or other readout. This technique enabled the identification of novel types of ipRGCs (Hatori et al., 2008; Ecker et al., 2010). Among retinal ganglion cells, the method appeared to selectively label ipRGCs. However, it also labeled other cells in the retina (rods and cones) and in the brain (e.g., cortical neurons) in which there is no known functional role for melanopsin. Some of this off-target labeling may result from transient developmental expression of melanopsin in tissues which don’t express it in adulthood; the early expression would trigger the recombination event, driving continuous reporter expression despite later downregulation of *opn4* gene expression. However, one study labeling using Cre-dependent viral vectors in adults, which circumvents the problem of transient developmental expression, nonetheless reported off-target labeling of a ganglion-cell type that lacks intrinsic photosensitivity (Delwig et al., 2016). It is unclear if melanopsin has other roles in these cells or whether the reporters used are so sensitive that they reveal even biologically insignificant levels of expression of the *opn4* gene.

These methods of labeling ipRGCs differ in which cell types they primarily reveal, as inferred from the morphology of labeled cells. Melanopsin immunostaining labels the outer “M1 plexus” of the IPL, corresponding to the dendrites of M1 ipRGCs as well as a fainter inner “M2 plexus” (Provencio et al., 2002; Berson et al., 2010). Other ipRGC types costratify in these bands, with the M3 and M6 cell stratifying in both bands (Berson et al., 2010; Quattrochi et al., 2018) and the M4 and M5 ipRGCs stratifying in or near the M2 band (Estevez et al., 2012; Stabio et al., 2017).

These differing methods also label different projections in the brain. The first genetic reporter labeled mainly M1 cells and revealed output projections to the intergeniculate leaflet (IGL), suprachiasmatic nucleus (SCN), olivary pretectal nucleus (OPN), and ventral lateral geniculate nucleus (vLGN) (Hattar et al., 2002, 2006). Cre-based reporters showed additional labeling in the ventromedial portion of the dorsal lateral geniculate nucleus (dLGN), and the stratum opticum of the superior colliculus (SC) (Brown et al., 2010; Ecker et al., 2010). Retrograde tracing has showed that these additional targets receive input from M4, M5, and M6 cells, all of which can generate an intrinsic photocurrent (Estevez et al., 2012; Stabio et al., 2017; Quattrochi et al., 2018).

Thus, there are a great variety of approaches for tagging ipRGCs, each with clear advantages, but each, also, with obvious shortcomings. Here we used morphology, retinal stratification, brain projections and abundance of labeled cells to evaluate the efficacy and specificity of six different methods of labeling ipRGCs. We show that two popular red-fluorescent Cre-reporter lines, Ai9 and Ai14, crossed with Opn4^cre^ mice, evoke off-target labeling of many conventional (non-ipRGC) ganglion cells. Additionally, we show that other Cre-reporter lines that label ipRGCs specifically underreport the total number of ipRGCs. Finally, by combining these methods with melanopsin immunostaining and Opn4-BAC transgenic reporters to label ipRGCs with multiple methods in the same mouse, we provide evidence that ipRGCs comprise a higher percentage of RGCs in the mouse retina than previously recognized.

## Methods

### Retinal Dissections

Retinal dissections were performed 3-6 weeks after viral injections. Both eyes were removed immediately after death and immersed in cold Ames medium (Sigma-Aldrich, St. Louis, MO). The retinas were carefully dissected from the eyecup by cutting around the cornea then removing the vitreous and lens. Four incisions were made to flatten and mount the tissues onto filter paper. The retinas were fixed for 30 minutes in 4% paraformaldehyde (Electron Microscopy Sciences, Hatfield, PA, 32%) prepared in 0.1M phosphate buffered saline (PBS; pH 7.4), then washed in PBS (3 x 15 minutes) before immunolabeling.

### Image Analysis & Acquisition

Epifluorescence images were acquired on a Nikon Eclipse microscope (Micro Video Instruments, Avon, MA) equipped with a SPOT RT Slider digital microscope camera (Diagnostic Instruments, Sterling Heights, MI), as described previously (Ecker et al., 2010). Composite images from single focal planes were stitched together in Adobe Photoshop.

Confocal images of tissues at different depths (“z-stacks”) were obtained on Zeiss LSM510 and LSM 800 Meta laser scanning confocal fluorescent microscopes. These were analyzed with Zeiss Zen software, Image J / FIJI and Adobe Photoshop. All image adjustments were global. Cell-count analyses were performed using Image J and R. Counts of cell densities were calculated based on labeled confocal stacks taken from regions at least 30% and no more than 70% of the distance from optic disc and retinal margin. Cells were marked as labeled only if they could be conclusively identified based on single channel labeling to avoid bias between different co-labeling experiments.

### Animals

All experiments were conducted strictly in accordance with the National Institutes of Health guidelines, and all laboratory animal use and care protocols were approved by the Institutional Animal Care and Use Committee at Brown University.

Mice (n=20) were obtained by mating homozygous Opn4^cre/cre^ mice (Ecker et al., 2010) with one of three lines of heterozygous transgenic Cre-reporter mice: “Z/EG” (Jackson Laboratories #00390, Bar Harbor, ME), “Ai9” (Jackson Laboratories #007909); “Ai14” (Jackson Laboratories #007914) as described previously. Opn4-GFP mice, a BAC transgenic strain generated by the GENSAT project (Gong et al., 2003), were obtained from MMRRC (033064-UCD), and crossed to Opn4^cre/cre^ animals.

Animals were maintained in individual cages in a 12:12 hour light:dark cycle. Mice were sacrificed between 6 weeks and 3 months of age by CO_2_ asphyxiation with immediate decapitation or lethal injection of Beuthanasia.

### Electrophysiology

Voltage-clamp recordings were conducted as described by (Estevez et al., 2012). Briefly, internal solutions consisted of 120 mM Cs-methanesulfonate with 5mM NaCl, 4mM CsCl, 2mM EGTA, 10mM HEPES, 4mM ATP-Mg, 7mM phosphocreatine-Tris, 0.3mM GTP-Tris, along with 2mM QX314 (to eliminate sodium spikes). The internal solutions were adjusted to pH 7.3. Osmolarity of the internal solutions was between 270–280 mOsm. To block synaptic transmission onto ganglion cells, we bath-applied a drug cocktail (Wong et al., 2005, 2007; Estevez et al., 2012) consisting of 100 μM L-(+)-2-amino-4-phosphonobutyric acid (L-AP4, a group III metabotropic glutamate receptor agonist), 40 μM 6,7-dinitroquinoxaline-2,3-dione (DNQX, AMPA/kainate receptor antagonist), and 30 μM D- (-)-2-amino-5-phosphonopentanoic acid (DAP5, NMDA receptor antagonist). Recordings were obtained with a Multiclamp 700A amplifier coupled to a Digidata 1322 digitizer with pClamp9.2 data acquisition software (Molecular Devices). Sampling frequency was 10 kHz with recordings low-pass filtered at 4 kHz. We did not compensate for series resistance; data were discarded if the series resistance exceeded 30 MΩ. Cells were voltage clamped at −64 mV after a liquid junction potential correction of −6.6 mV in our system. Pipettes for voltage-clamp experiments were pulled from a Flaming/Brown P97 pipette puller (Sutter Instruments) with tip resistances ranging from 4–8 MΩ when submerged in the bath.

### Brain Histology

Two to six weeks after intraocular injections, mice were sacrificed by CO_2_ asphyxiation and cervical dislocation. Brains were removed and immersed in 4% paraformaldehyde for 24 hours. On the following day, the brain was transferred to phosphate-buffered saline and molded in a 4% agarose gel. Brains were sectioned coronally at 50 μm on a vibratome (Leica VT1000S). Serial sections spanning all targets of optic axons, from the optic chiasm to the superior colliculus, were mounted on glass slides or processed for immunohistochemical labeling.

### Ocular Injections

Mice of age ~P14-45 were anesthetized by continuous inhalation of isoflurane (2% in oxygen). Respiration rate and isoflurane delivery were monitored throughout the surgery. Under a dissecting scope, tweezers were used to apply gentle pressure on either side of the eye to protrude it and a small volume of axon-tracing reagent was injected through the sclera using a micropipette. We used two such reagents. The first was Cholera toxin β-subunit conjugated to Alexa-594 (5μg/μL, approx 0.5μL). The second was a Cre-dependent viral tracer that expresses a red fluorescent protein in transduced neurons (DIO-eF1α-ChR2-mCherry AAV2/2; University of North Carolina Vector core, [Chapel Hill, NC), titer: 10^12^ viral molecules per ml). Postsurgical care included analgesia (buprenorphine SR; 0.1 mL/g subcutaneously) and topical ophthalmic ointment. Animals were monitored for recovery daily for 2 days following the surgery and euthanized 2-6 weeks later for tissue harvest.

### Immunohistochemistry

Brain slices were placed into a blocking solution of 10% goat serum and 0.25% Triton X in phosphate-buffered solution [PBS) for 2 hours. Primary antibody was added and the tissue kept cold overnight (4 deg. C). After a wash (3 x 10 minutes in PBS), the tissue was immersed in the secondary antibody solution for 2 hours, washed as before, and mounted on glass slides.

Retinas were fixed for 30 minutes in 4% paraformaldehyde and rinsed in PBS. Melanopsin labeling was amplified with tyramide-signaling amplification (TSA) using horseradish peroxidase (HRP)-tagged goat anti-rabbit IgG and Alexa Fluor 594 tyramide (TSA-15, Molecular Probes, Eugene, OR). The protocol in the kit was followed except that we replaced amplification buffer with PerkinElmer 1X Plus Amplification Diluent. GFP signal was amplified using anti-GFP primary antibodies and tdTomato signal was amplified anti-mCherry antibodies (See Table 1). Incubation times were 2-3 days for the primary antibody and overnight for the secondary antibody, both at 4°C. Retinas were mounted on glass slides and cover-slipped with Aqua-Mount (Invitrogen, Carlsbad, CA).

**Table 1:**
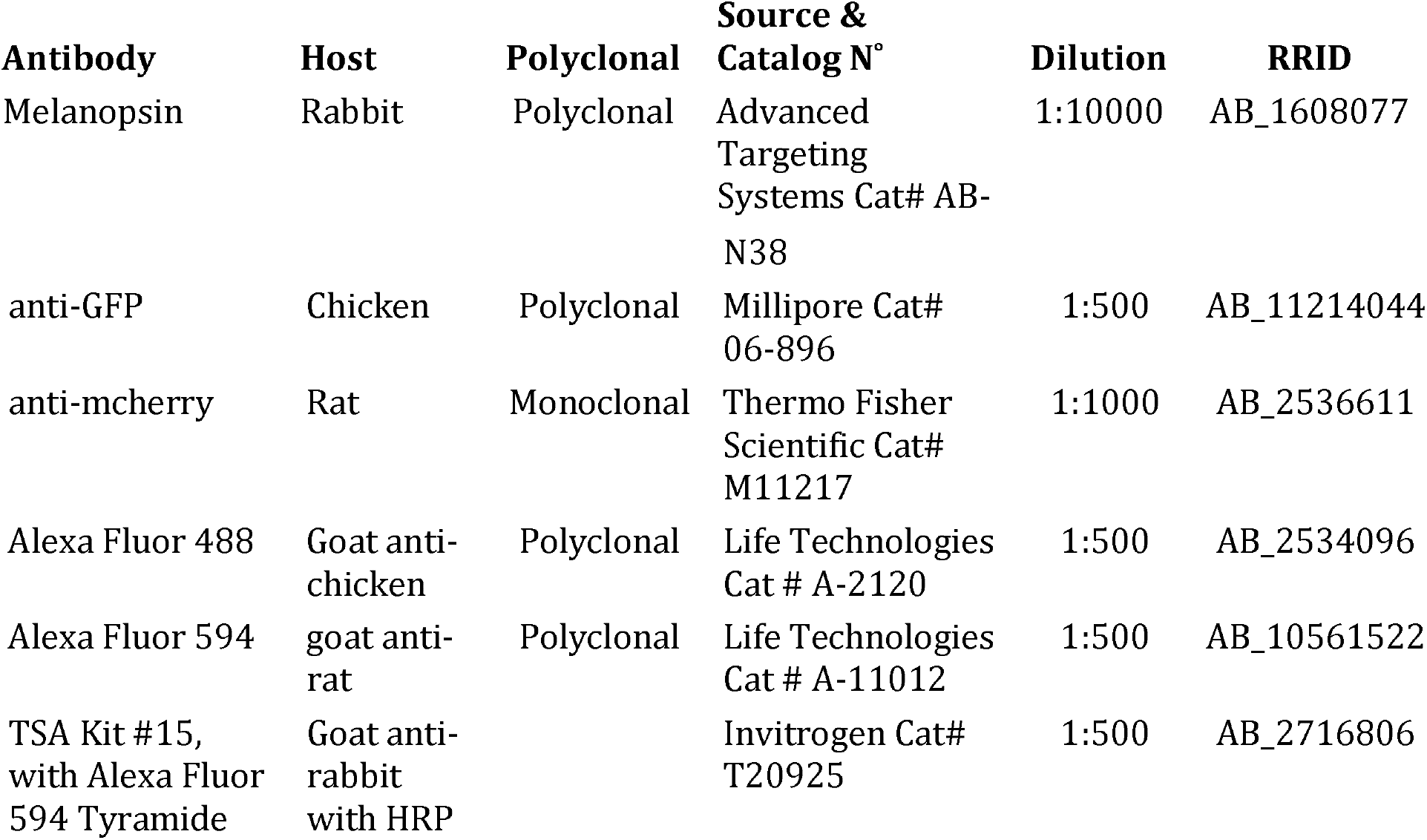
Antibodies used.

## Results

To attempt to capture the entire population of ipRGCs, we turned to the most sensitive reporter system known to selectively label them. Based on available literature, this is the Opn4^cre/+^; Z/EG^+/-^ line, which has been shown to label six types of ipRGCs (M1-M6) (Ecker et al., 2010; Stabio et al., 2017; Quattrochi et al., 2018) By contrast, melanopsin immunostaining labels only M1-M3 cells reliably, although weak somatic labeling of M4, M5 and M6 cells can sometimes be detected (Berson et al., 2010)

Despite its selectivity, we find this Cre-based system is surprisingly inefficient at labeling ipRGCs. Nearly half of melanopsin-immunopositive ipRGCs (i.e., presumptive M1-M3 cells) lacked detectable GFP-immunofluorescence in Opn4^cre/+^; Z/EG^+/-^ retinas (Figure 2) (41%; 95% binomial confidence interval 37-66%). Reversing this comparison, we find that only about a third of GFP-labeled cells in Opn4^cre/+^; Z/EG^+/-^ retinas were melanopsin immunoreactive (35%; 95% binomial confidence interval 32-38%). This is expected because this Cre-based system is known to reveal ipRGC types that lack detectable immunoreactivity (Ecker et al., 2010).

**Figure 1:**
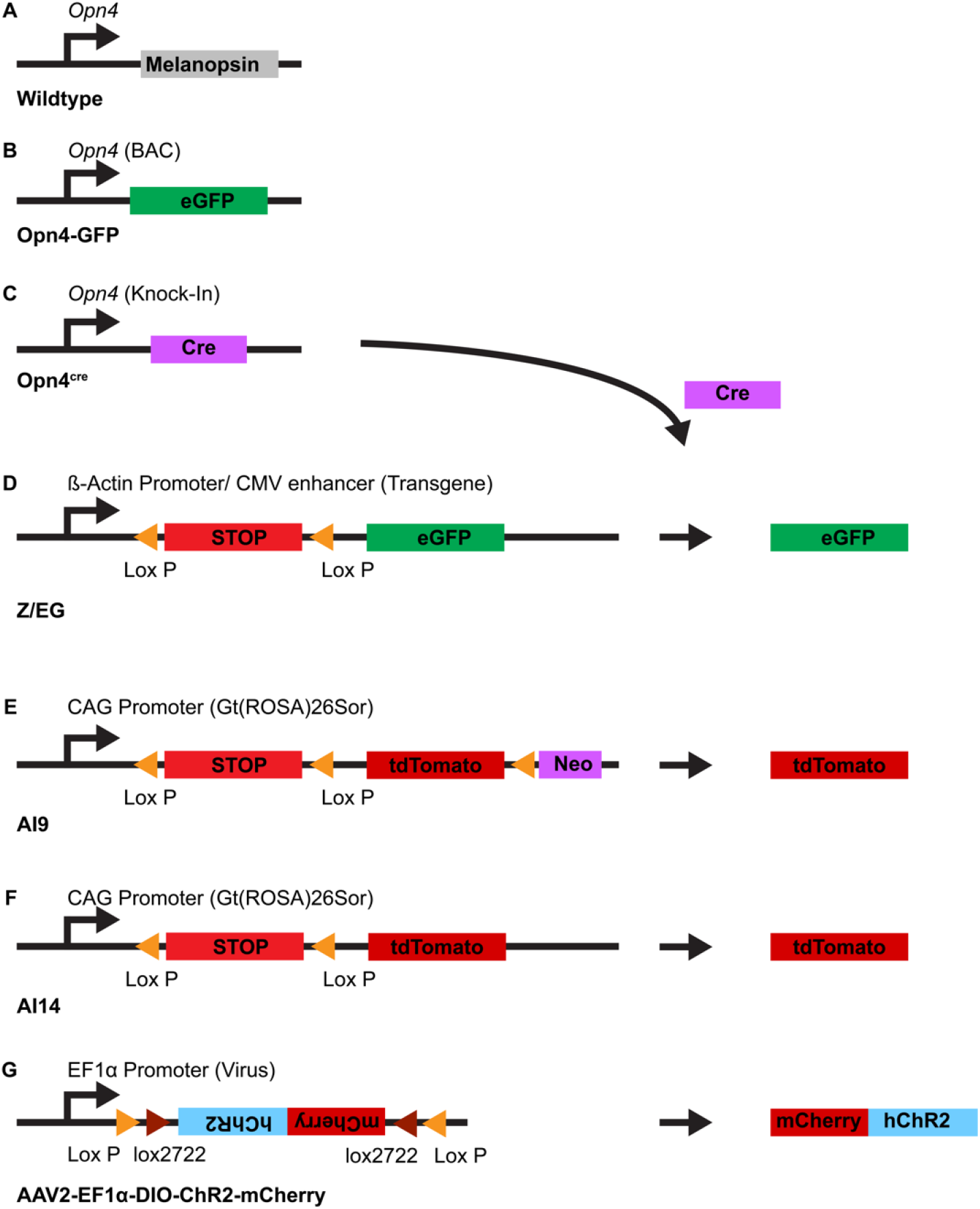
Mouse genetic models for labeling ipRGCs. A: In wildtype mice, expression of melanopsin is under control of the opn4 gene. B: Opn4-GFP mice have been genetically modified by a transgene delivered by a bacterial artificial chromosome in which the melanopsin gene has been replaced with one coding for the fluorescent reporter eGFP. **C:** Opn4^cre^ mice have the the cre gene knocked into the opn4 gene locus so that Cre is expressed under the control of the melanopsin promoter. Cre can trigger permanent recombination in a number of reporters, leading to persistent expression of the Cre-dependent reporter through the lifetime of the animal. **D-G:** Cre-dependent reporters: **D:** The Z/EG reporter expresses eGFP after recombination under control of the B-Actin promoter. E: The Ai9 reporter, knocked into the highly efficient Rosa 26 locus, expresses tdTomato after recombination under control of the CAG promoter. F: The Ai14 reporter is a variant of the Ai9 reporter with the neomycin cassette removed, leading to less labeling in the absence of Cre. G: AAV2-EF1a-DIO-ChR2-mCherry labels infected Cre-expressing cells with the ChR2-mCherry fusion protein, which is localized to the membrane of neurons.

**Figure 2:**
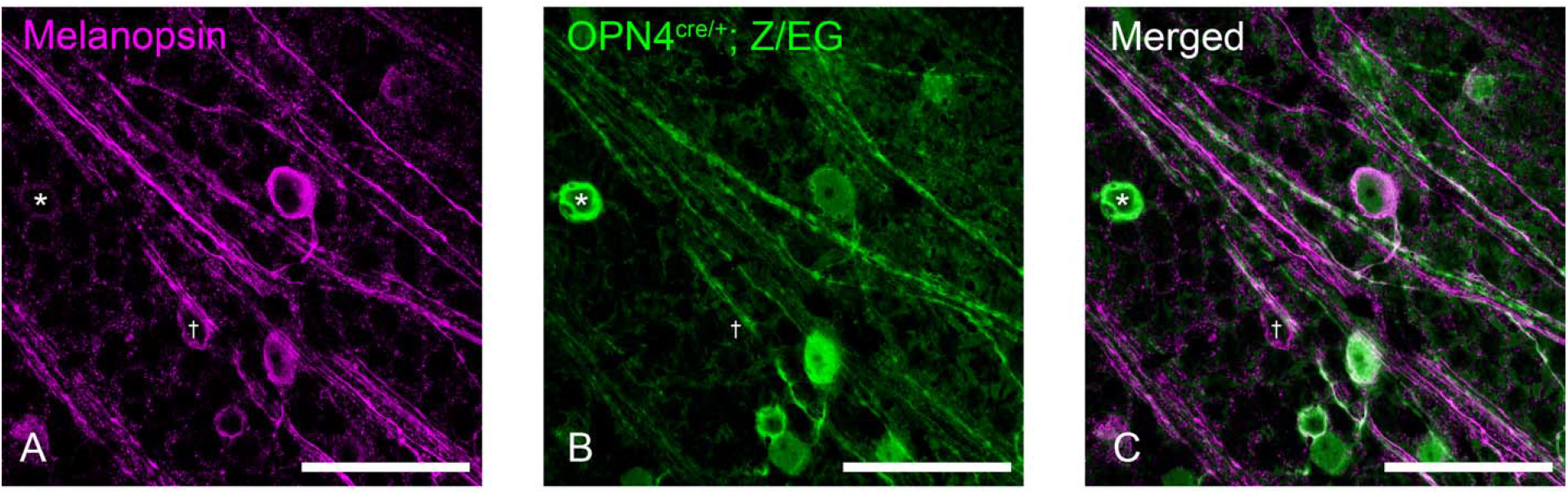
Confocal images of melanopsin and anti-GFP immunostaining of retinal wholemounts in Opn4^cre/+^; Z/EG animals. A: Anti-melanopsin TSA-enhanced staining of ganglion cell layer. B: GFP labeling of cell bodies and axons reveal by anti-GFP immunofluorescence. C: Merged. Note presence of cells labeled by GFP but not melanopsin (*), and cell that is melanopsin immunolabeled but lacks anti-GFP staining (†). Scalebar 100μm.

Spurred by this evidence that both this genetic approach and conventional immunolabeling substantially underreport the full population of ipRGCs, we sought to enhance the efficiency of genetic labeling of ipRGCs by applying various more sensitive Cre reporters in Opn4^cre^ mice. In one approach, we evoked red fluorescent labeling in Cre-expressing ganglion cells by injecting an AAV2 virus carrying a Cre-dependent mCherry construct. In other studies, we evoked red fluorescence instead by crossing Opn4^cre/cre^ mice with two strains of highly efficient Cre reporter mice (Ai9 and Ai14; tdTomato reporter). An additional technical advantage of these methods is that unlike those previously considered they also label the efferent axons of labeled RGCs. This allowed us to assess patterns of axonal termination in the retinorecipient visual nuclei as well as the labeling of ipRGCs within the retina.

### Cre-dependent viral labeling

We injected the eyes of Opn4^cre/+^ mice with a Cre-dependent virus (AAV2.2 DIO-eF1α-ChR2-mCherry; Fig 3). This induced red fluorescent labeling of a mixture of RGCs. Many corresponded to M1-M3 ipRGCs because they were melanopsin immunopositive. Others exhibited little or no demonstrable immunoreactivity (Fig. 3A), but the evidence suggests most if not all of these were also ipRGCs, belonging to types M4, M5 and M6, which exhibit little immunoreactivity for melanopsin. Their dendrites were appropriately restricted to the melanopsin immunoreactive plexuses, as is true of all known ipRGCs whether immunoreactive or not (Berson et al., 2010; Ecker et al., 2010; Estevez et al., 2012; Stabio et al., 2017; Quattrochi et al., 2018). Further, their labeled axons terminated only in retinorecipient targets known to receive ipRGC input (Fig. 4), including the suprachiasmatic nucleus, olivary pretectal nucleus, ventral lateral geniculate nucleus, intergeniculate leaflet, stratum opticum of the superior colliculus and the ventromedial segment of the dLGN (Fig 4) (Hatori et al., 2008; Ecker et al., 2010; Delwig et al., 2016). The accessory optic nuclei were free of labeled optic fibers, as expected for selective labeling of ipRGCs, because these nuclei are selectively innervated by direction-selective ganglion cells which are not ipRGCs. This contrasts with the off-target labeling of axons in the MTN, an accessory optic nucleus, in an earlier study that used a particularly sensitive Cre-reporter virus in Opn4^cre^ mice (Delwig et al., 2016). The virus used in the present study thus appears to report melanopsin expression more faithfully.

**Figure 3:**
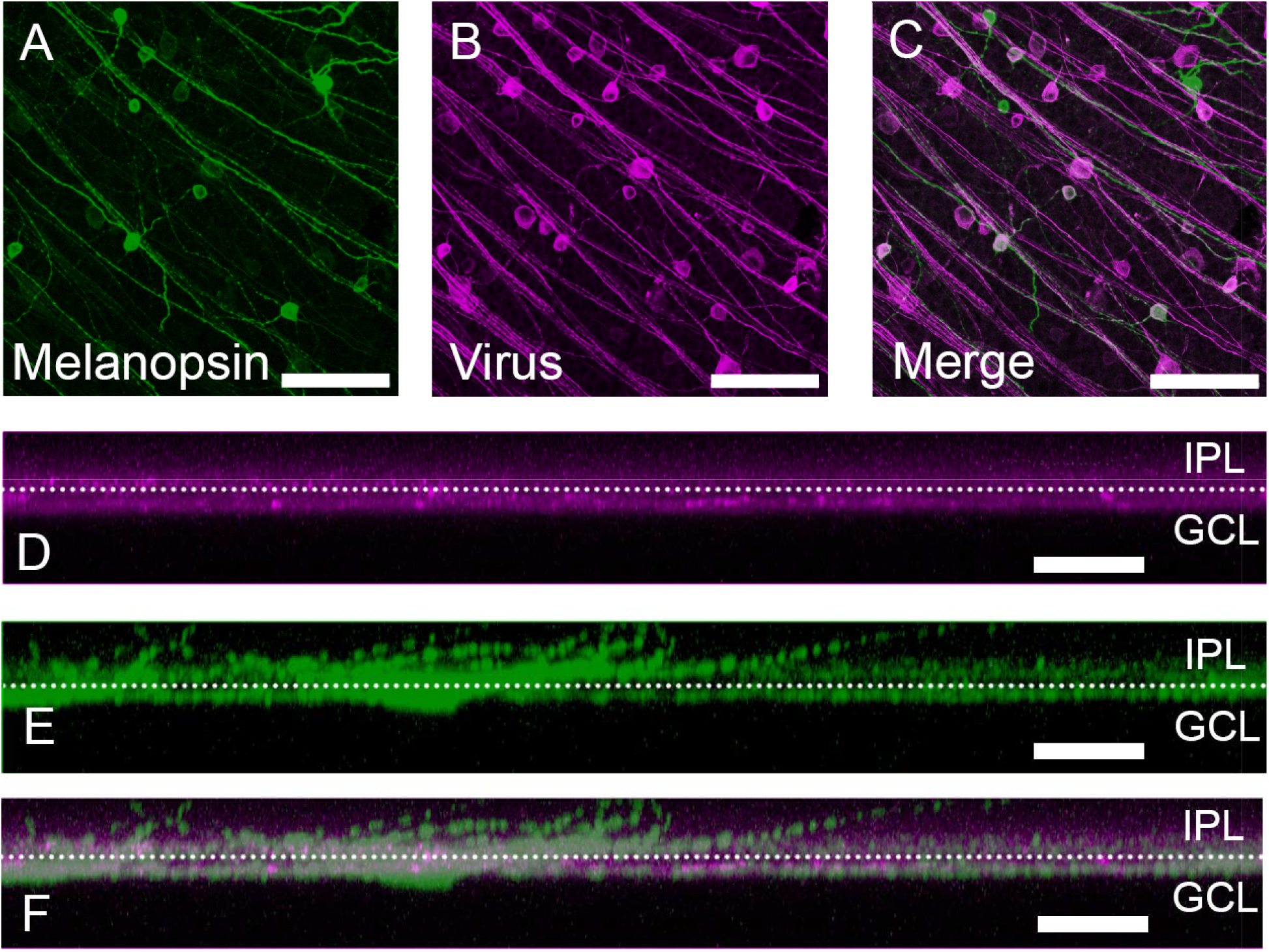
Confocal images of ganglion-cell layer in wholemount retina of anti-melanopsin and anti-mCherry immunolabeling in Opn4^cre^ animals infected with a Cre-dependent virus (AAV2-EF1-DIO-ChR2-mCherry). A: Melanopsin TSA immunolabeling of a single optical slice through the ganglion-cell layer. **B:** Anti-mCherry immunostaining of the same field of view. **C:** Merged. D-F: Side-view projections showing stratification of labeled processes in the opticfiber layer, ganglion-cell layer and inner part of the inner plexiform layer of retina. D: Melanopsin TSA immunolabeling. E: Anti-mCherry labeling F: Merged. Scalebar 100μm (A-C), 20μm (D-F).

**Figure 4:**
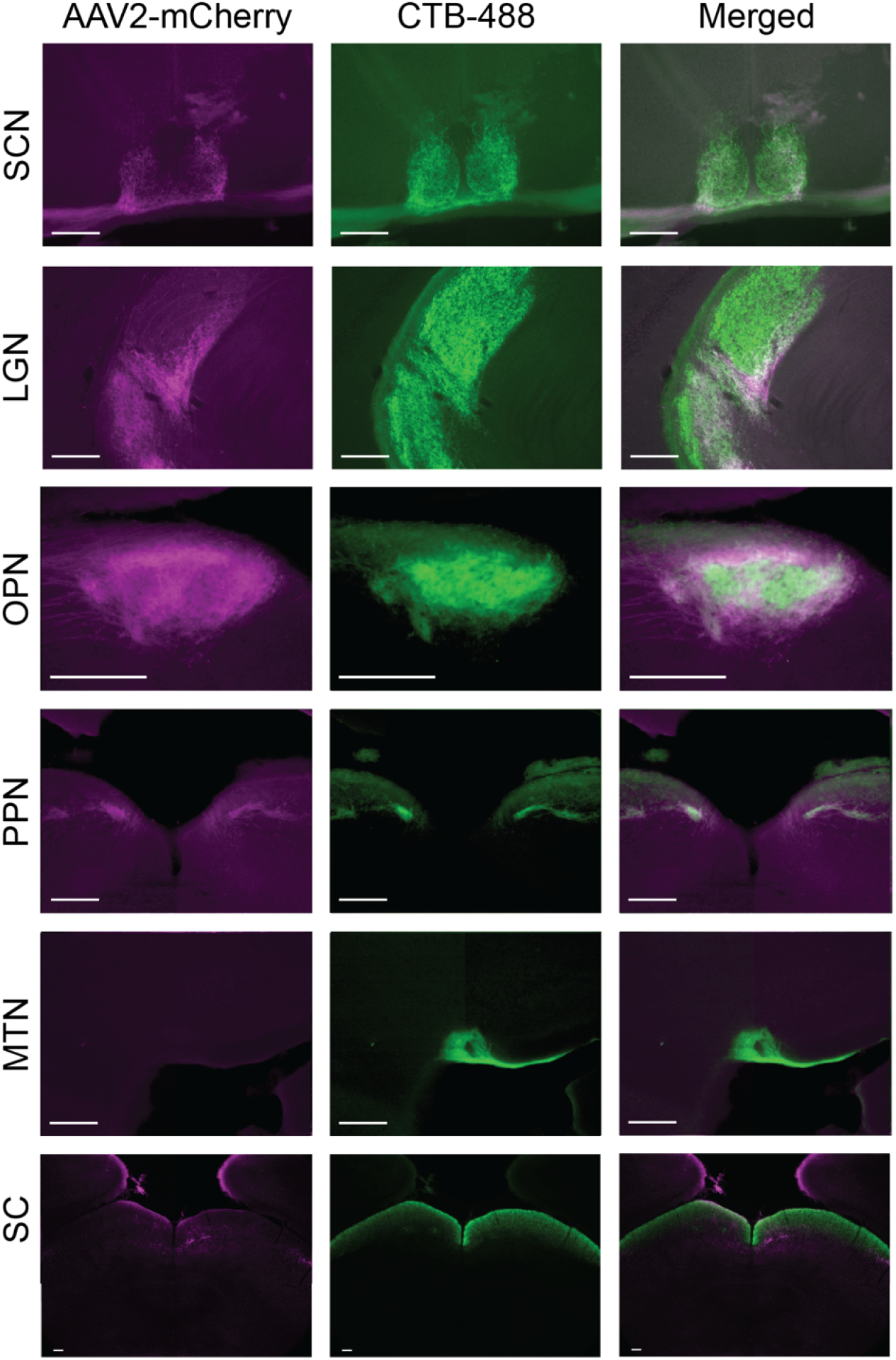
Labeling of retinal targets in Opn4^cre/+^ mice injected intraocularly with a Cre-dependent virus (AAV2-EF1a-DIO-ChR2-mCherry) and with the anterograde tracer CTB-488. Fluorescent images of viral labeling (left), CTB-488 (middle) and merged (right) in coronal sections containing the suprachiasmatic nucleus (SCN), lateral geniculate nucleus (LGN), olivary pretectal nucleus (OPN), posterior pretectal nucleus (PPN), medial terminal nucleus (MTN), and superior colliculus (SC). Virally labeled projections project to a subset of ocular targets marked by CTB-488, including the SCN, vLGN, ĨGL, and central dLGN, the OPN, PPN, and stratum opticum of the superior colliculus, but not the MTN. Scalebar 100μm.

The density of cell bodies labeled by the virus (180 cells/mm^2^) was similar to that obtained with the Opn4^cre/+^; Z/EG reporter system (177 cells/mm^2^), which our data show significantly underreports the full population of ipRGCs. In the case of the viral labeling, this underreporting derives at least in part from the low efficiency of the virus in transducing the M1 ipRGC type, presumably due to viral serotype. Because M1 cells express the highest levels of melanopsin and the strongest melanopsin immunoreactivity among ipRGCs, we could use the immunofluorescence to trace their dendrites to their arborization in the outermost IPL. Most of these dendrites and their parent cell bodies lacked viral labeling. In a single sample area with robust viral transduction, we counted a total of 39 identifiable M1s. Less than a third of these (30%; 12/39) of were virally labeled.

In contrast, nearly every immunolabeled M2 cell was well labeled by the virus. In the same sample region, 96% of identifiable M2 cells were virally labeled (26/27). Virally labeled M4 (ON alpha) cells were also readily identifiable from their large cell bodies and thick dendrites. Smaller virally labeled RGCs with little or no melanopsin staining presumably represent mainly M5 and M6 ipRGCs. The successful transduction of M6 cells is supported by the presence of a fine plexus of virally labeled dendrites in the outermost IPL. These were immunonegative for melanopsin, and thus presumably not dendrites of M1 or M3 cells, the only other ipRGC types known to stratify at this level.

### More sensitive Cre-reporter mouse strains suffer from off-target labeling

Because none of the methods considered to this point fully labeled the ipRGC population, we asked whether we might do better with more efficient Cre reporters. We first crossed Opn4^cre/cre^ mice to the sensitive Ai9 Cre reporter strain (Madisen et al., 2009) which drives robust tdTomato labeling through use of the highly efficient Rosa26 locus and CAG promoter. This Opn4^cre/+^; Ai9 cross clearly augmented the labeling, yielding a density of labeled RGCs more than three times higher than that in the Opn4^cre/+^; Z/EG cross considered previously (Ai9: 583 cells/mm^2^; Z/EG: 176 cells/mm^2^) (Figure 5, see also Figure 10). However, the pattern of labeling suggested than many RGC types known or believed to lack intrinsic photosensitivity were among the labeled population. Dendritic labeling spanned essentially the full thickness of the IPL rather than being restricted to the known ipRGC strata. Cells with morphology clearly distinct from the known ipRGC types were labeled (Figure 5D), and recordings from a small sample of these confirmed that they lacked intrinsic photosensitivity (Figure 5E). We conclude that in this context, as in others (Madisen et al., 2009), the Ai9 line allows ‘leaky’ expression of the reporter in cells that do not express Cre. We therefore turned to the Ai14 reporter (Fig 5 F-H) a modification of the Ai9 line that reportedly shows less of this off-target labeling (Madisen et al., 2009). In our hands, however, Opn4^cre/+^; Ai14 mice produced a pattern of retinal labeling similar to that in Opn4^cre/+^; Ai9 mice. The density of labeled ganglion cells was only slightly lower (517 cells/mm^2^) than in the Ai9 cross. Both Opn4^cre/+^; Ai9 (Figure 6) and Opn4^cre/+^; Ai14 mice (Figure 7) exhibited apparent off-target labeling of optic fibers in the medial terminal nucleus and the outer shell of the dLGN, which are known to lack ipRGC input. The full distribution of labeled optic axons could not be determined because of masking by widespread reporter expression in brain neurons (Ecker et al., 2010)

**Figure 5:**
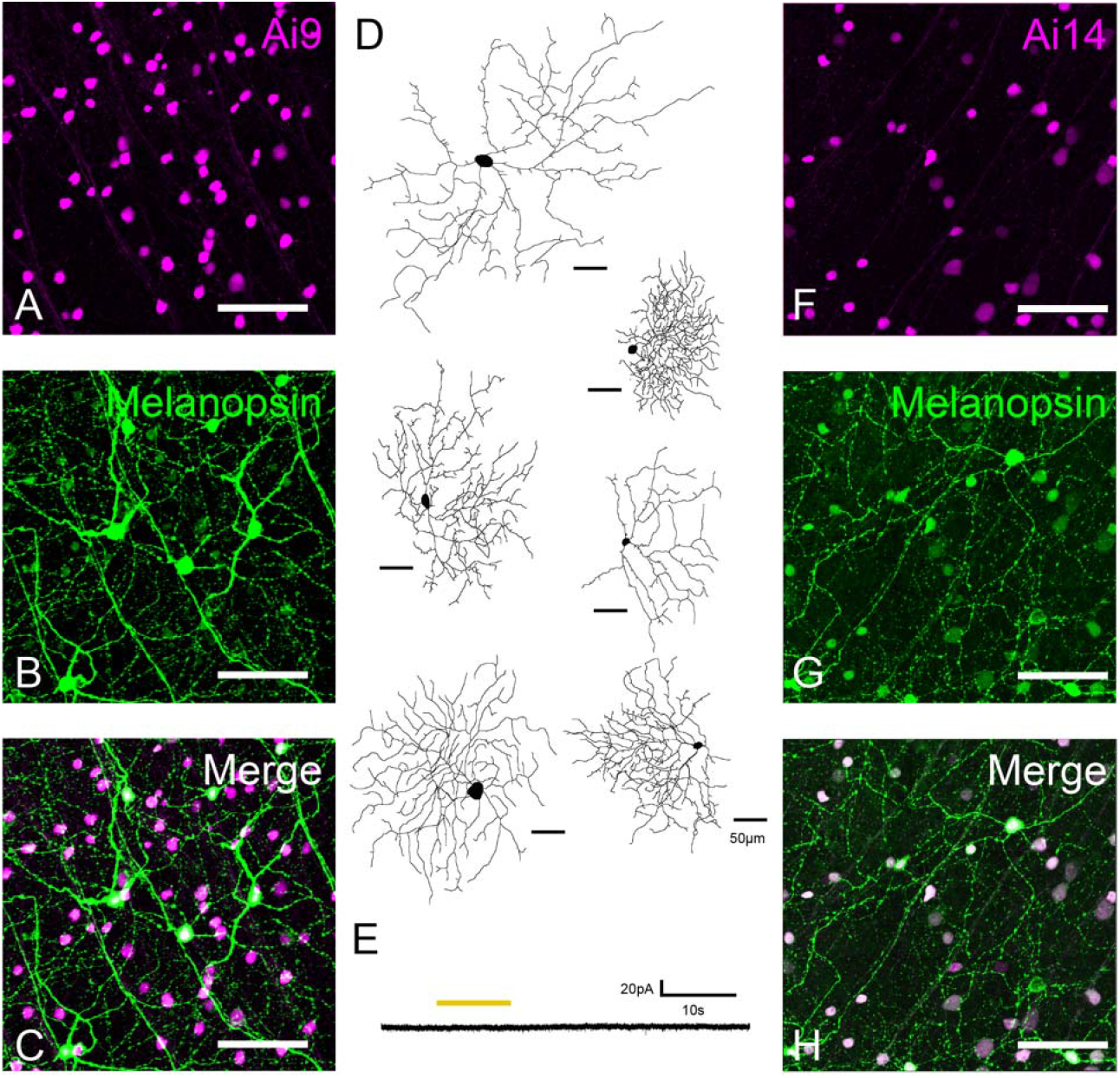
Retinal ganglion cells labeled by crossing the Opn4^cre/+^ mice with two strains of Cre-reporter mice: Ai9 and Ai14. **A:** Confocal images of tdTomato-labeled somas in the ganglion cell layer of Opn4^cre/+^; Ai9 mice. B: Anti-melanopsin TSA immunostaining in the same field of view. C: Merged image of A and B. D: Reconstructions of Opn4^cre/+^; Ai9-tdtomato labeled retinal ganglion cells showing diversity of labeled cell types E: Absence of intrinsic light response in an RGC labeled in an Opn4^cre^; Ai9-tdTomato mouse; voltage-clamp recording (V_hold_ −64 mV) during pharmacological block of rod and cone input. Yellow bar indicates stimulus timing. **F-H:** As in A-F, except using the Ai14 reporter. Scalebars are 50 ¼m in D and 100μm in A-C and F-H.

**Figure 6:**
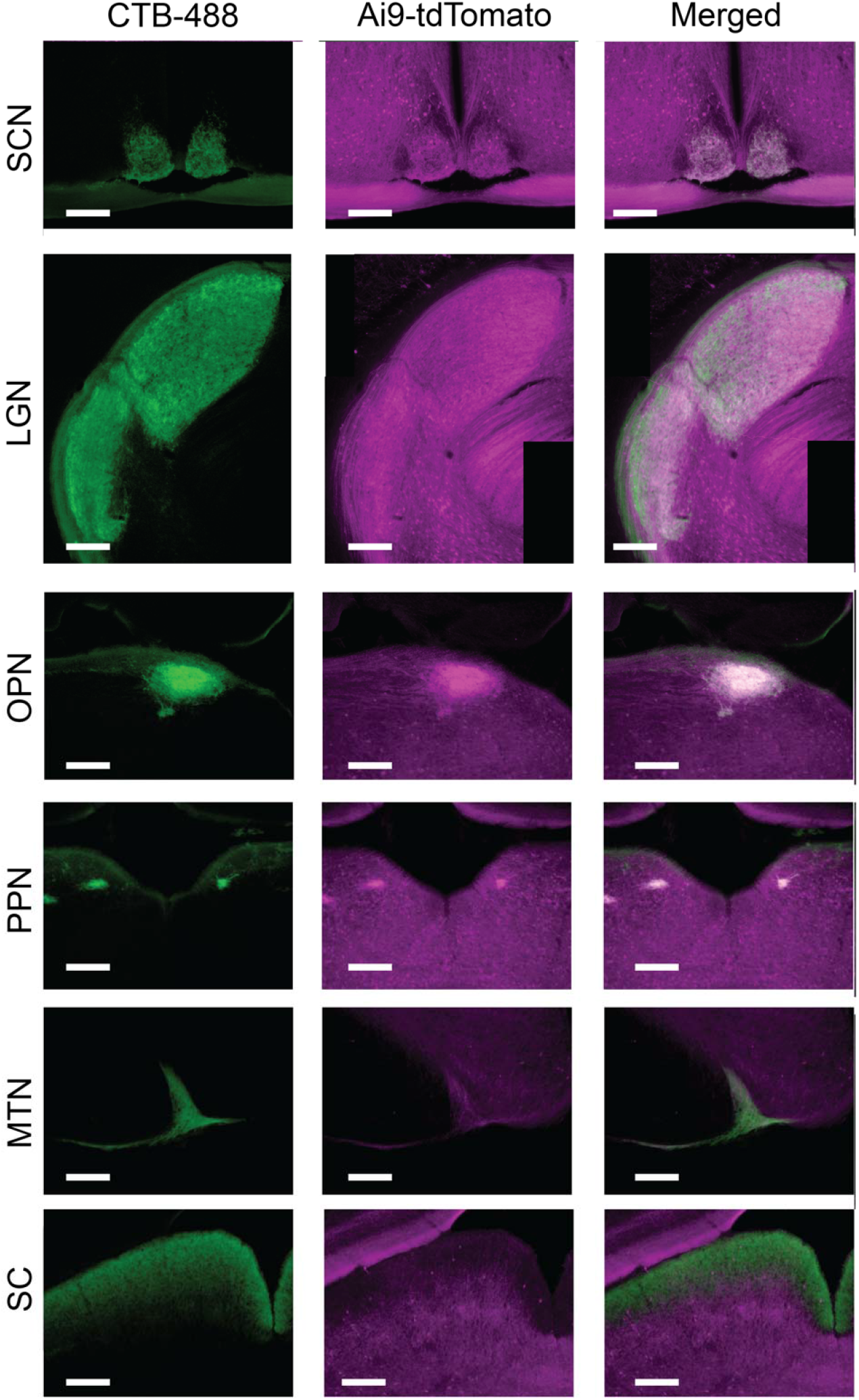
Anterograde labeling of optic fibers in various retinal targets in Opn4^cre/+^; Ai9 mice after intraocular injection of CTB-488. Fluorescent images of CTB-488 labeling (left), tdTomato labeling (middle) and merged (right) in coronal sections containing the suprachiasmatic nucleus (SCN), lateral geniculate nucleus (LGN), olivary pretectal nucleus (OPN), posterior pretectal nucleus (PPN), medial terminal nucleus (MTN), and superior colliculus (SC). In addition to labeling fibers in the outer shell of the dLGN, which is thought to lack ipRGC input, tdTomato expression in Opn4^cre/+^; Ai9 mice labels neuronal cell bodies in the brain. Scalebar 100μm.

**Figure 7:**
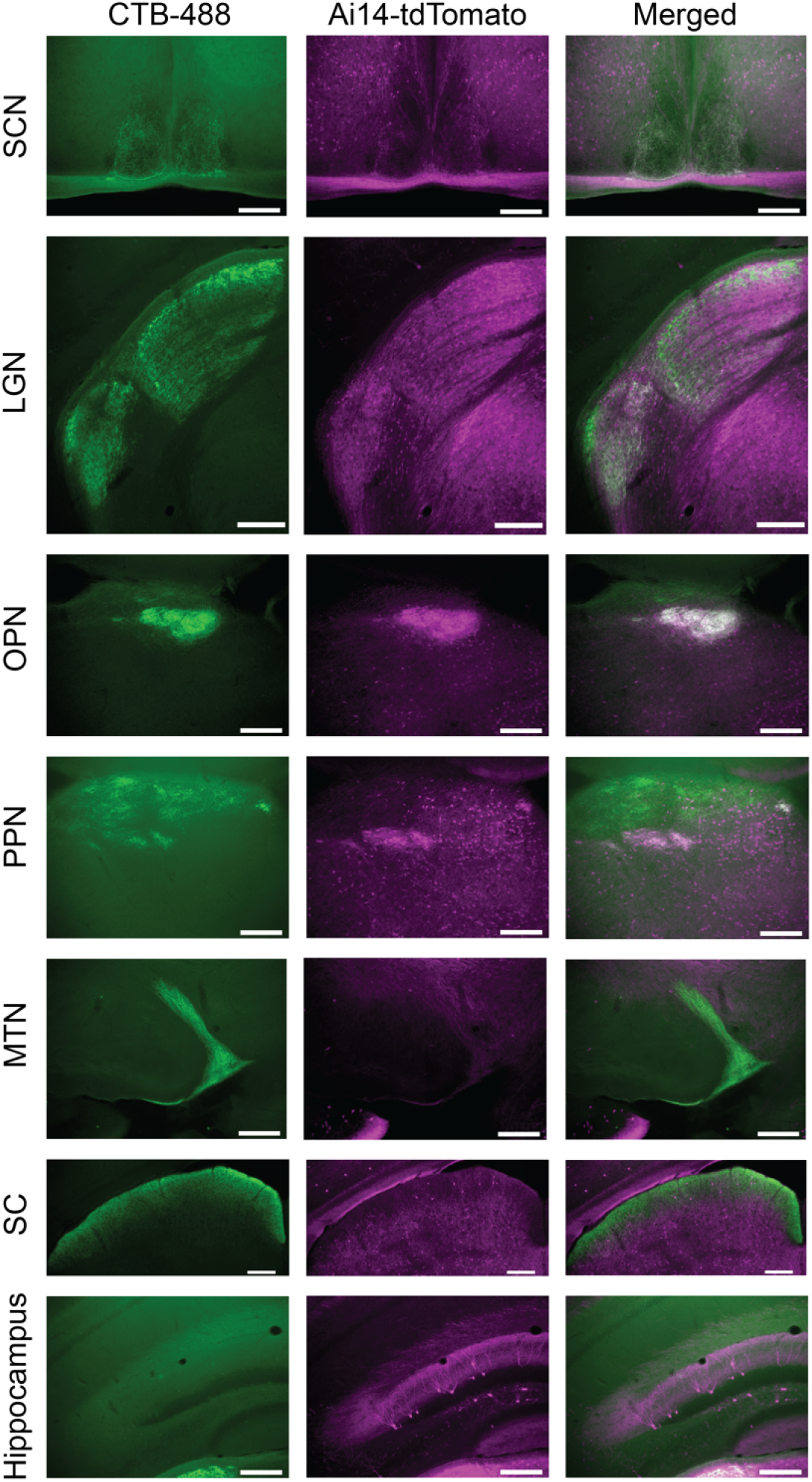
Anterograde labeling of retinal targets in Opn4^cre/+^; Ai14 mice after intraocular injection of CTB-488. Fluorescent images of CTB-488 labeling (left), tdTomato labeling (middle) and merged (right) in coronal sections containing the suprachiasmatic nucleus (SCN), lateral geniculate nucleus (LGN), olivary pretectal nucleus (OPN), posterior pretectal nucleus (PPN), medial terminal nucleus (MTN), superior colliculus (SC), and hippocampus. tdTomato labeling is evident in the outer shell of the dLGN, as well as in cell bodies in the brain, including the hippocampus. Scalebar 100μm.

### A novel transgenic melanopsin reporter is selective but incomplete

In the face of the shortcomings of the Cre-based approaches, we vetted one additional genetic reporter, a BAC transgenic mouse line developed by the GENSAT project for the melanopsin gene, here termed Opn4-GFP.’ The brain projections suggested this reporter might be labeling mainly M1 cells because they closely matched those in the Opn4^tau-LacZ^ mice, in which mostly M1 cells appear to be labeled (Hattar et al., 2006). Labeling was largely restricted to the SCN, OPN shell, PPN, IGL and, more faintly, the vLGN. The MTN and SC were essentially devoid of labeling (Figure 9).

Preferential labeling of M1 cells would not be surprising for a transgene expressed under the regulation of the melanopsin promoter because M1 cells express the highest levels of melanopsin of any ipRGC type. In fact, though, we saw labeling of many M2 cells along with M1 cells, and found that many cells of both types were unlabeled. GFP was detectable in more than two thirds of M1 cells (69%; 27/39) and more than three quarters of M2 cells (77%; 21/27), identified as above by the stratification of their immunolabeled dendrites. The density of labeled cells in Opn4-GFP retinas (69 cells/mm^2^) was substantially lower than that revealed by melanopsin immunostaining (103 cells/mm^2^) (Fig 8, see also Fig 10). Slightly less than half of melanopsin immunopositive cells were labeled by GFP (43%, 95% binomial confidence interval 38-48%), while just over half of the GFP+ cells exhibited melanopsin immunolabeling (58%, 95% binomial confidence interval 52-63%). Many of the GFP+ neurons that lacked melanopsin immunoreactivity had small somas which may indicate off-target labeling of some amacrine cells.

**Figure 8:**
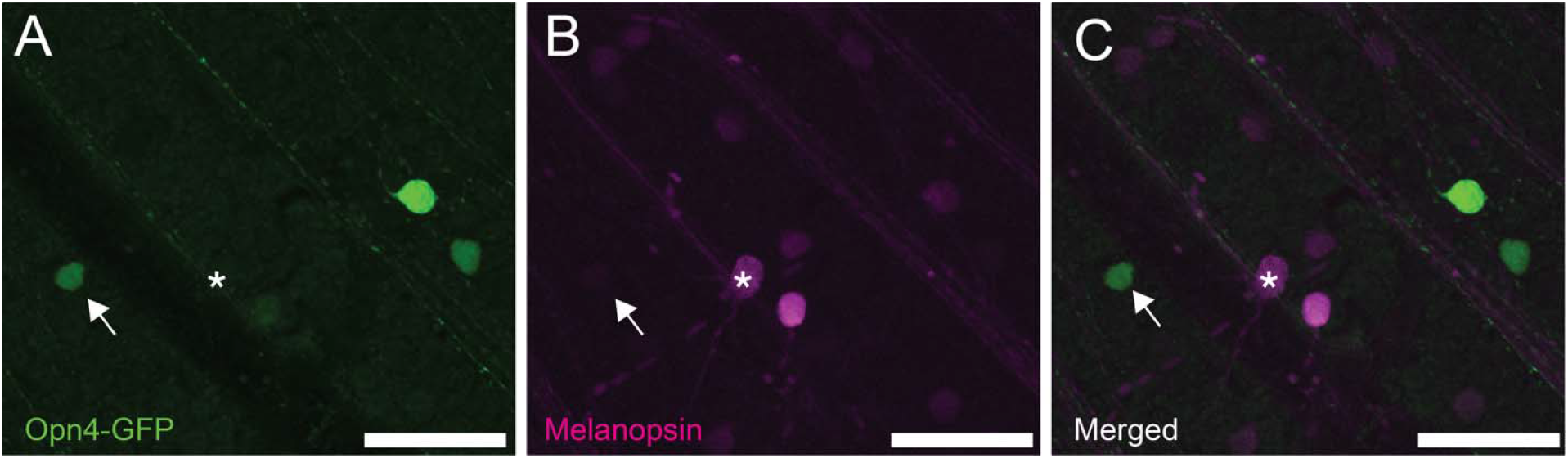
Confocal images of the ganglion cell layer in a retinal flatmount from an Opn4-GFP mouse, comparing the labeling patterns produced by the transgene (A) and melanopsin immunofluorescence (B). C: Merged. Arrow indicates a GFP-labeled cell without appreciable melanopsin immunostaining, while * represents a cell with melanopsin immunostaining but no transgene expression. Scalebars 50μm

**Figure 9:**
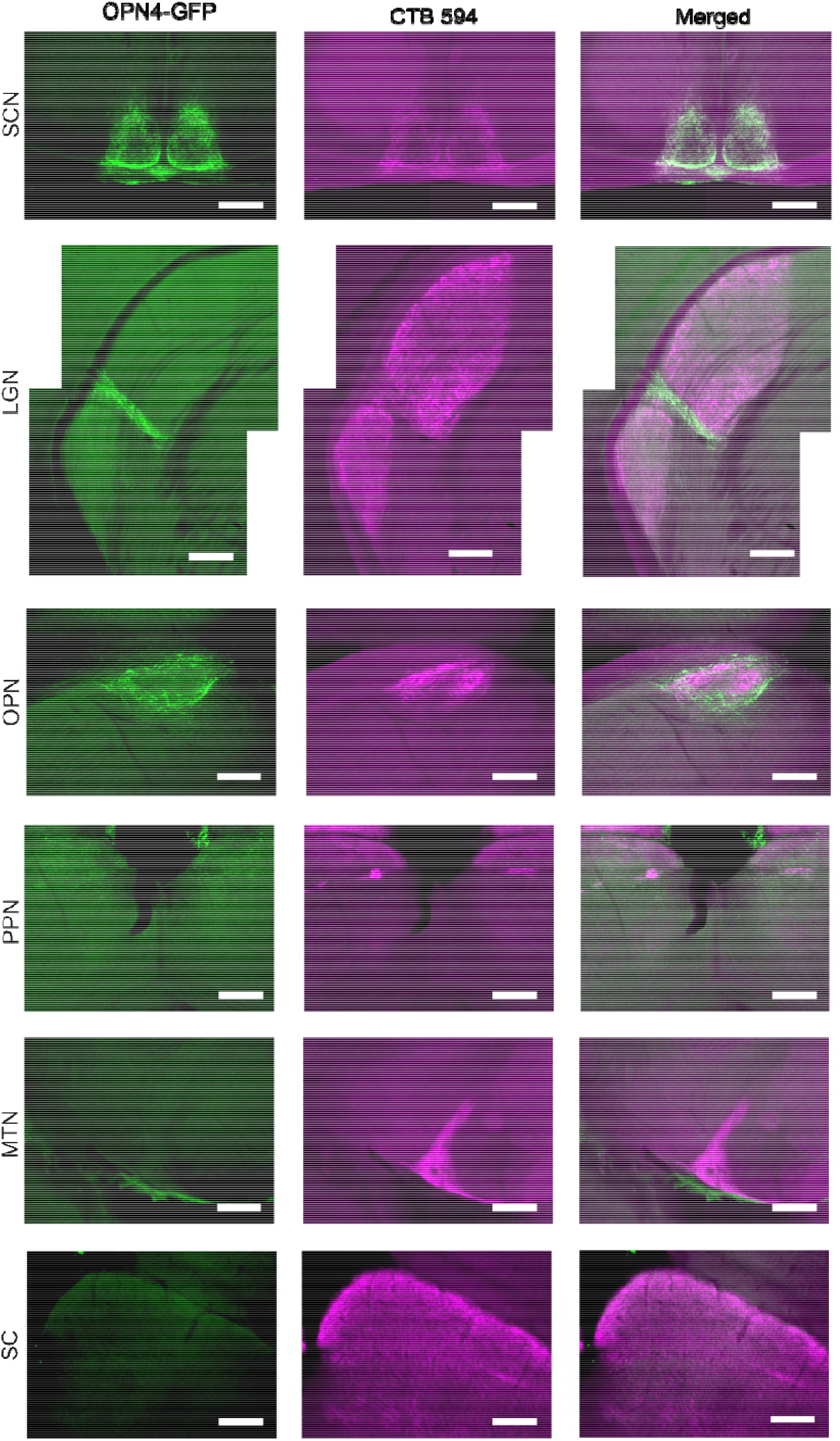
Labeling of retinal targets in Opn4-GFP mice by intraocular injections of CTB-594. Fluorescent images of anti-GFP immunostaining (left), CTB-594 labeling (middle), and merged (right) in coronal sections containing the suprachiasmatic nucleus (SCN), lateral geniculate nucleus (LGN), olivary pretectal nucleus (OPN), posterior pretectal nucleus (PPN), medial terminal nucleus (MTN), and superior colliculus (SC). GFP labeling is evident in the SCN, IGL, vLGN, and shell of the OPN. Scalebar 100μm.

**Figure 10:**
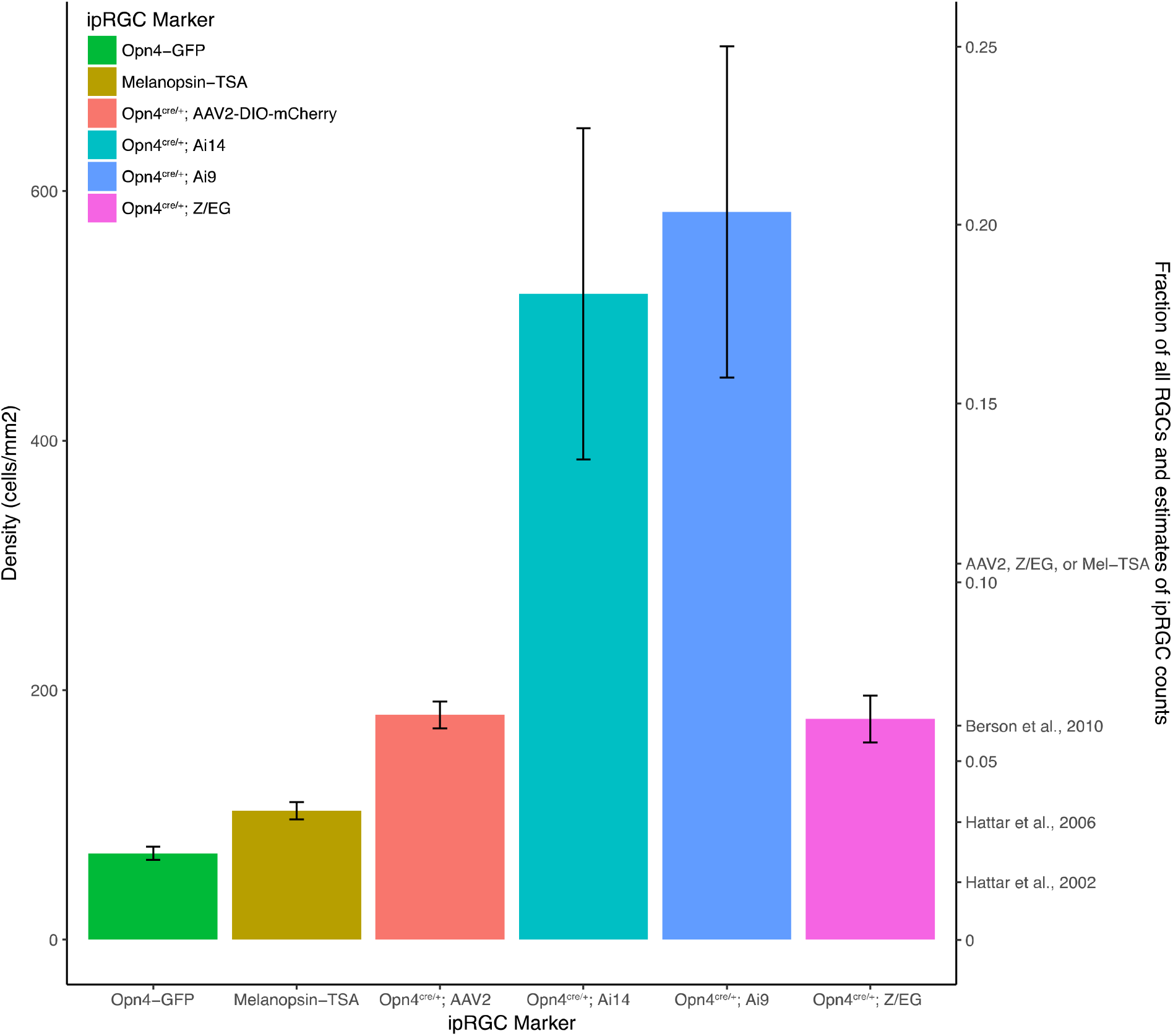
Density of labeled cells for different putative methods of labeling ipRGCs. **Left Axis:** Density of retinal ganglion cells labeled by different methods of labeling ipRGCs. **Right Axis:** Estimate of total percentage of RGCs labeled by each method, as well as previous estimates of the total number of ipRGCs in the mouse retina. Using multiple faithful methods in combination (AAV2, Z/EG, and Melanopsin TSA) reveals that the total number of ipRGCs may be higher than previously reported. Error bars: SEM.

### Quantitative Comparisons of Methods for Labeling ipRGCs

To this point, we have established that none of the various reporters we have assessed labels all ipRGCs and only ipRGCs. To provide an overview of the relative completeness of these reporter systems and to reassess the global abundance of ipRGCs, we compared the density of labeled somas across reporters, where possible within the same animals. These values were converted to percentages of all RGCs based on previous estimates of the total area of the mouse retina (15.6mm^2^) and the total number of RGCs (44,860) (Jeon et al., 1998)

None of the three ipRGC-selective markers labeled more than 7% ganglion cells overall. Melanopsin immunolabeling, which labels all M1-M3 cells but omits the M4-M6 types, labeled 4%. The Opn4-GFP transgenic mouse, again labeling mostly M1 and M2 cells, marked only 2% of all ganglion cells. That value is almost certainly inflated by inclusion of GFP-positive displaced amacrine cells and, for RGCs, is consistent with selective but incomplete labeling of M1 and M2 cells. The two faithful variations of the Cre-based methods, the Z/EG reporter cross and AAV2 viral labeling, each labeled close to 7% of all ganglion cells. Crosses of the Opn4^cre^ mouse with the sensitive reporter strains (Ai9; Ai14) labeled many more RGCs, but apparently mainly through extensive off-target labeling.

The actual fraction of ipRGCs must be higher than the lower bound of 7% established by the best methods used isolation because we have shown that none of these methods labels all ipRGCs. To assess the degree of redundancy among these faithful markers, we conducted two sets of triple-labeling experiments.

In the first set of experiments, we injected Cre-dependent AAV expressing mCherry into Opn4^cre/+^; Opn4-GFP mice while in the second set of experiments injected the same virus into Opn4^cre/+^; Z/EG mice. In both cases we immunostained against melanopsin, mCherry, and GFP and compared the incidence of co-labeling among the different markers. Some cells, especially M2 cells, were labeled by all three reporters in these experiments, but many others exhibited only one or two of them (Figure 11). Overall, about a tenth of all RGC somas were labeled by at least one of AAV2, Z/EG, or melanopsin immunostaining, suggesting 10% as a more accurate lower bound for the proportion of ganglion cells that are intrinsically photosensitive in mice.

**Figure 11:**
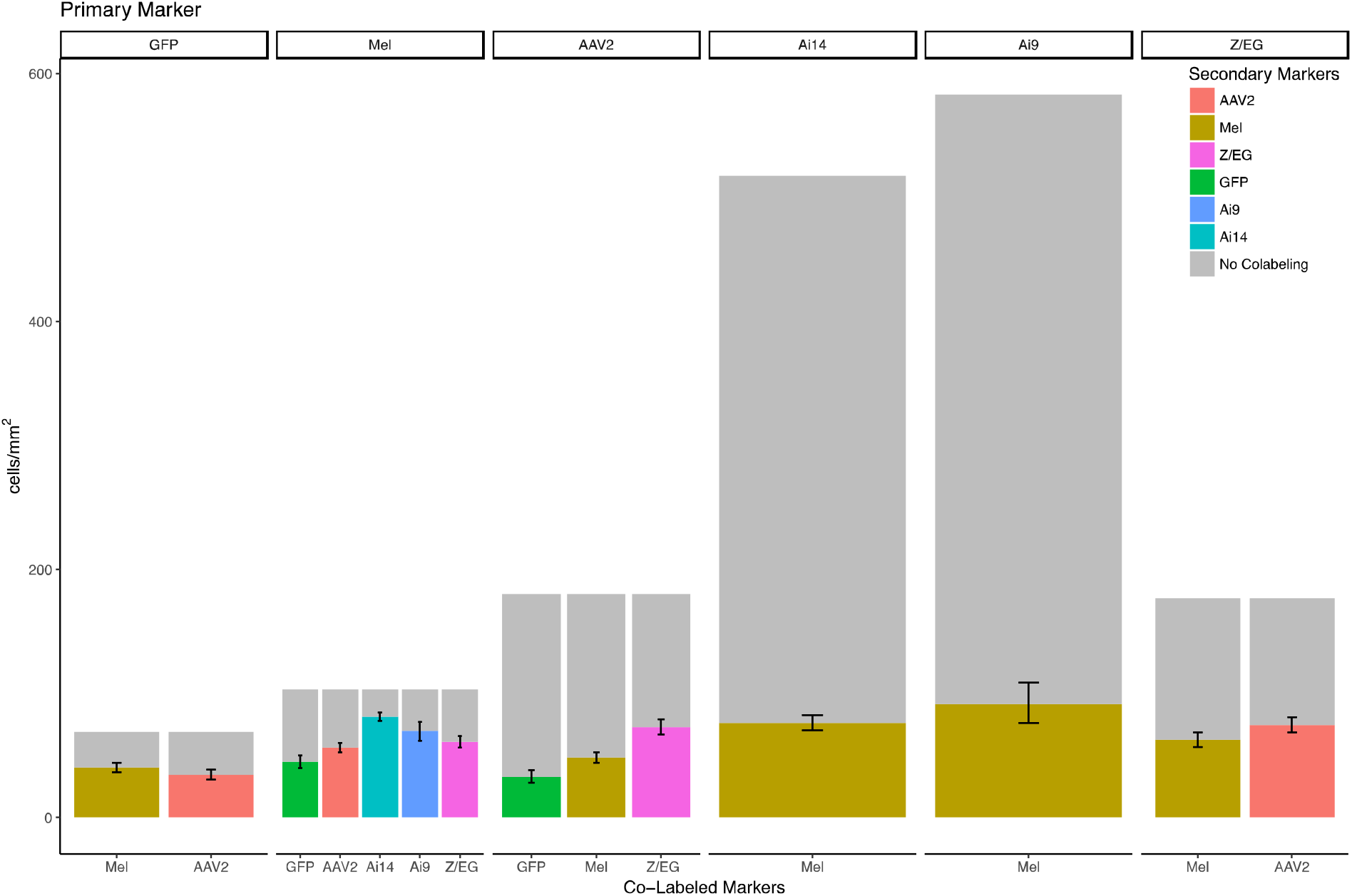
Co-labeling among ipRGC markers. For each ipRGC primary marker (top), the graph indicates how much of the population of labeled cells is also labeled by other markers. Error bars: 95% binomial confidence intervals.

## Discussion

We have surveyed the merits and limitations of diverse methods for labeling ipRGCs in mice. Some approaches proved of little value. Others that are widely used suffer from various shortcomings that may limit their utility in specific experimental settings. Together, our findings demonstrate the importance of matching the labeling approach used the experimental question being addressed, of acknowledging the limits of the method used, and of documenting the selectivity and efficacy of any novel reporter system.

### The Opn4^cre^ melanopsin-driver line can trigger off-target labeling in retina and brain

Both antibody staining and transgenic reporters for *opn4* expression reveal ipRGC types that express high levels of melanopsin (M1, M2 and M3 cells), but the amplification offered by Cre technology has proven essential for revealing the types with lower expression: M4 (ON alpha), M5 (Pix ON), and M6 types (Hatori et al., 2008; Brown et al., 2010; Ecker et al., 2010)

This greater efficiency comes at a cost, though. Our results using highly amplifying Cre reporter strains (Ai9 and Ai14) reinforce earlier evidence that the most widely used Opn4^cre^ mouse strain can trigger off-target labeling of neurons in both retina and brain (Ecker et al., 2010; Delwig et al., 2016). The amount of off-target labeling depends on the Cre reporter. The sensitive Cre-dependent viral vector used by Delwig (Delwig et al., 2016) appeared to label ipRGCs with reasonable specificity, but it also exhibited some off target labeling of the rare ON direction-selective cells. By contrast, our crosses with two sensitive Cre-reporter mouse strains produced extensive off-target labeling. About a fifth of all ganglion cells were labeled, including many that were clearly not ipRGCs, as reflected by their dendritic morphology, a lack of intrinsic photosensitivity, and off-target labeling of retinal axons in inappropriate brain targets.

Fortunately, two methods seem to allow the Opn4^cre^ line to drive labeling exclusively in melanopsin-expressing RGCs and not in other RGC types. First, the Opn4^cre/+^;; Z/EG cross apparently labels only ganglion cells that express melanopsin, and no other RGCs, though it does suffer from off-target labeling of outer retinal photoreceptors and brain neurons. The second successful approach was intraocular injection of a Cre-dependent mCherry viral vector in Opn4^cre^ mice. This appeared to label large numbers of ipRGCs of several types, including at least M1, M2, M4 and M6 cells. Brain projections included all the targets identified in studies using the original knock-in reporter (Opn4^tau-lacZ^) which appears to label M1 ipRGCs in relative isolation, despite the fact that the virus we used transduces M1 ipRGCs with low efficiency. Labeled axons also terminated in some targets not revealed by the LacZ reporter, consistent with its robust labeling of other ipRGC types, especially the M2 cell. Significantly, the virus we used did not transduce ON direction-selective ganglion cells because a target of such cells, the MTN, contained no labeled optic axons. This contrasts with clear MTN labeling an earlier study using a different virus in the same melanopsin-driver mouse (Delwig et al., 2016). This may be due to a difference in viral serotype since we used an AAV2.2 whereas the virus in the other study was an AAV2.1. Serotype interactions with cell-type-specific molecular features also may also explain why only about a third of M1 cells were labeled by the virus we used, contrasting with the nearly ubiquitous labeling of M2 cells. Differences between the promoter used in the viral construct we exploited (EF-⍰) from the CMV promoter widely used in other Cre-dependent viruses might play a role as well (Zhang et al., 2010; Ho et al., 2015).

A particular advantage of the viral approach is that it allows Cre-mediated recombination to be restricted in space (eye only vs. whole animal) and in time (postnatally, to avoid labeling cells that only express melanopsin transiently during development). However, off-target labeling can clearly occur even with such temporal control (Delwig et al., 2016). This Cre system thus can produce false-positive labeling in adult retinas due to shortcomings beyond the transient developmental expression of melanopsin.

### Popular faithful methods for labeling ipRGCs are incomplete and label disparate mixtures of cell types

We show that two widely used methods for labeling ipRGCs ⍰ melanopsin antibodies and GFP tagging in Opn4^cre/+^; Z/EG mice ⍰ label partially overlapping subsets of ipRGCs. Though both methods mark only cells with functional melanopsin expression, neither marks all such cells.

These results are consistent with the limitations of melanopsin immunostaining, which depends upon strongly expressed melanopsin and thus fails to label M4 (ON alpha), M5 (Pix ON) and M6 cells, despite functional evidence that those cell types exhibit melanopsin-dependent intrinsic photocurrents (Ecker et al., 2010) On the other hand, the genetic approach left half of melanopsin immunopositive M1-M3 cells unlabeled by GFP. Thus, although the genetic labeling method is far more effective than immunolabeling for revealing ipRGC subtypes that weakly express melanopsin (M4-M6), its labeling of ipRGCs is nonetheless very incomplete.

Our work highlights the importance of validating individual Cre-reporters when exploiting Opn4^cre^ mice because Cre appears to be expressed in many RGCs lacking both melanopsin immunoreactivity and intrinsic photosensitivity. Of course, new *bona fide* ipRGC types may yet emerge with more ideal reporters or other methods. Further, the opn4-driven Cre expression in diverse non-photosensitive neurons in both retina and brain may yet prove to be functionally meaningful because melanopsin expression may contribute to non-photosensory cellular functions.

### Available methods underrepresent the true abundance of ipRGCs

The faithful ipRGC-labeling methods provide some leverage for assessing the densities and relative abundance of ipRGCs among ganglion cells as a whole. However, because we conclude that no available strategy labels *all* and only ipRGCs, each method that labels ipRGCs selectively, can provide only a lower-bound estimate when used in isolation.

Overall, our estimates when using single labeling methods alone are in reasonable agreement with prior studies (Hattar et al., 2002, 2006; Berson et al., 2010). But here, by directly comparing several of these methods in single retinas, we show that each method labels a distinct subset of ipRGCs that partially but incompletely overlaps that labeled by other methods. Which ipRGCs are labeled is related in some cases to known distinctions among ipRGCs, such as cell type and level of melanopsin expression, but in other cases the basis for the incompleteness is unclear. When we accept any labeling of RGCs by any of the faithful labeling methods as evidence of ipRGC identity, we estimate that at least 11% of RGCs in mice express melanopsin and are capable of intrinsic phototransduction. This is higher than any other previous estimate, but it may still be an underestimate because some ipRGCs may have escaped detection by any of the methods used in combination in this analysis. One sanity check on this estimate comes from a comparison of the number of ipRGC types (5 or 6 depending on whether rare M3 variety is considered a true type) to the estimated total number of RGC types in mice (~40-50) (Baden et al., 2016; Bae et al., 2018; Goetz et al., 2022). By this measure, ipRGCs account for about 10-15% of all RGC types, close to the estimated percentage of all RGCs that are ipRGCs.

Our revised estimate comes with several potential caveats. In Opn4^cre/+^; Z/EG mice a small percentage of cells lack an intrinsic photocurrent (Ecker et al., 2010), raising the possibility that even relatively selective methods may label a few non-ipRGCs. The same might be true for at least some of the other apparently faithful labeling methods used in our analysis, which would leading us to overestimate the total abundance of ipRGCs. On the other hand, given the incomplete efficacy of any method for labeling ipRGCs, it is possible that a population of ipRGCs may be unlabeled by any of the methods described in this paper, either due to type specific biases or stochastic processes that cause these methods to label specific ipRGCs.

### Differing methods of labeling target different types of ipRGCs with differing efficiency

Given that none of the methods labels all members of all ipRGC types, the choice of which method to use when investigating ipRGCs will depend on the goal of the study and the desired target population, as each method labeled different populations in the eye and brain (Figure 12). Melanopsin immunostaining requires no special genetic manipulation, but labels primarily the highly expressing M1 and M2 ipRGCs. The Opn4-GFP line used here, like similar previously analyzed melanopsin transgenic reporter lines (Schmidt et al., 2008; Do et al., 2009), labels a subset of M1 and M2 neurons. These models, unlike *post hoc* immunolabeling, allow for targeting these ipRGC types *in vivo* and selective tracing of ipRGC projections to visual centers of the brain. On the other hand, the Opn4-GFP transgenic line we used labeled fewer true ipRGCs than any other method. The Opn4^cre^ mouse, in which *cre* has been knocked into the melanopsin gene locus, permits labeling or other manipulations of diverse ipRGC types, but it can both underreport and overreport the ipRGC population, depending on the Cre reporter system used. Some of the most sensitive Cre-reporter mouse lines should be avoided entirely because they produce off-target labeling in retina and brain. And both Cre reporter methods that avoided this problem (Opn4^cre/+^;; Z/EG mice and a flexed AAV2.2 vector) label only a subset of melanopsin immunopositive ipRGCs. The viral method is particularly inefficient for revealing M1 cells, presumably because of serotype specificity or factors linked to the promoter. The viral approach is particularly useful for limiting the time of Cre recombination to the age of injection, but if efficient labeling of M1 cells is needed, new variants of the AAV2 viral construct, using different combinations of promoter and serotype, should be considered.

**Figure 12:**
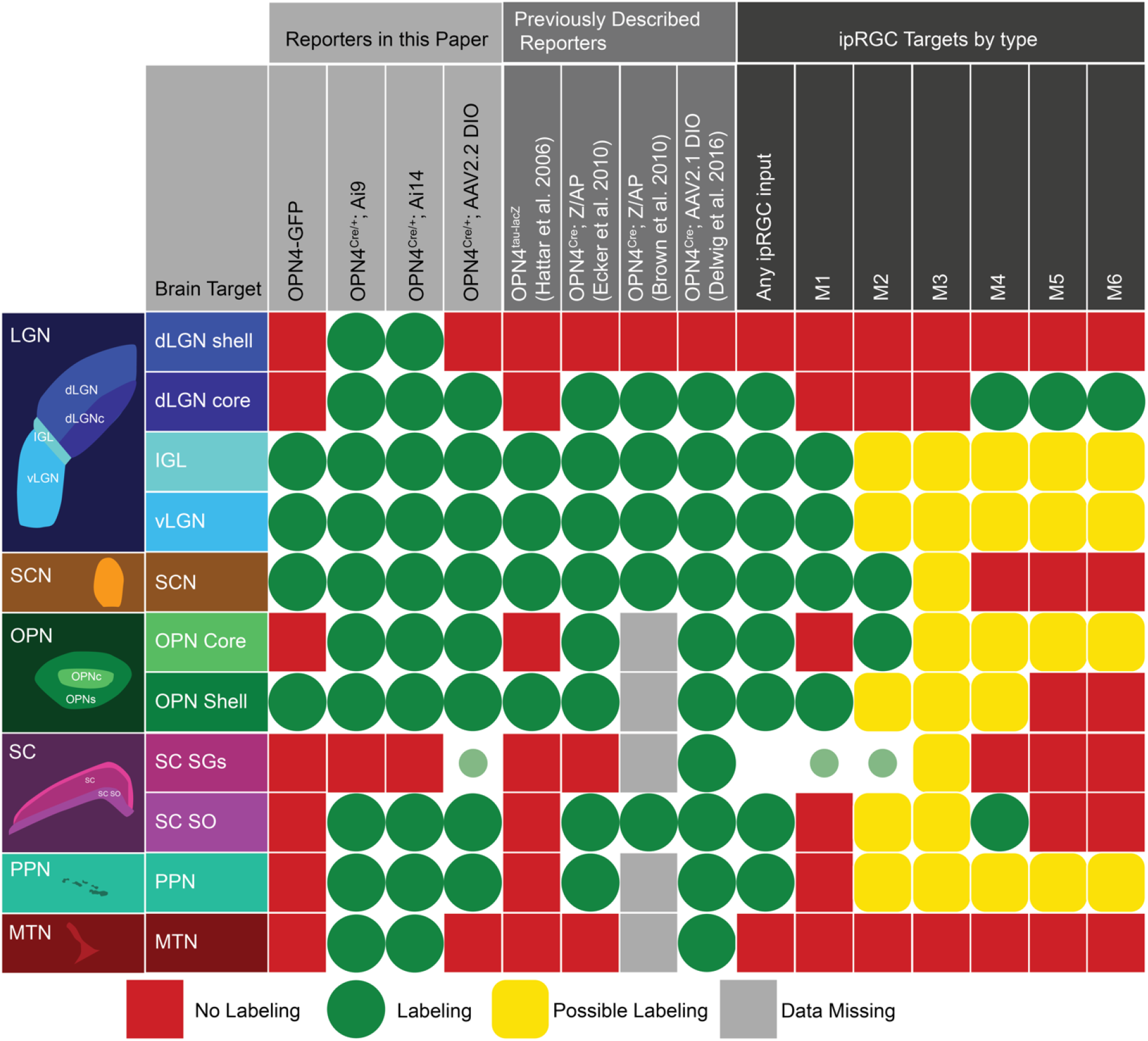
Schematic summary of patterns of optic-fiber labeling in major retinorecipient visual nuclei by various reporter systems (left half of grid). Reporter methods used in this paper appear in the leftmost columns, and those used in earlier studies appear to their right. For comparison, columns to the far right summarize the innervation of the same targets by individual ipRGC types (M1-M6) as inferred from previous studies.

